# Template switching mechanism drives the tandem amplification of chromosome 20q11.21 in human pluripotent stem cells

**DOI:** 10.1101/2020.07.11.198382

**Authors:** Jason A Halliwell, Duncan Baker, Kim Judge, Michael A Quail, Karen Oliver, Emma Betteridge, Jason Skelton, Peter W Andrews, Ivana Barbaric

## Abstract

Copy number variants (CNVs) are genomic rearrangements implicated in numerous congenital and acquired diseases, including cancer. In human pluripotent stem cells (PSC), the appearance of culture-acquired CNVs prompted concerns for their use in regenerative medicine applications. A particularly common problem in PSC is the occurrence of CNVs in the q11.21 region of chromosome 20. However, the exact mechanisms of origin of this amplicon remains elusive due to the difficulty in delineating its sequence and breakpoints. Here, we used long-range Nanopore sequencing on two examples of this CNV, present as a duplication in one and a triplication in another line. The CNVs were arranged in a head-to-tail orientation in both lines, with sequences of microhomologies flanking or overlapping both the proximal and distal breakpoints. These breakpoint signatures point to a specific mechanism of template switching in CNV formation, with surrounding *Alu* sequences likely contributing to the instability of this genomic region.

## Introduction

Copy number variants (CNVs) are gains or losses of DNA segments ranging in size from around 50bp to several megabases^1^. By affecting the dosage of genes and regulatory regions within the amplified or deleted sequence, CNVs underpin the aetiology of many diseases from developmental disorders to cancer^1^. The profound effect of the CNV acquisition on cellular phenotype has been also described in human pluripotent stem cells (PSC), which frequently gain a CNV located on chromosome 20 in the region q11.21 upon prolonged culture^2-5^. Once gained, the 20q11.21 CNV bestows on the variant PSC attributes that provide them with a growth advantage due to resistance to apoptosis^5,6^. The 20q11.21 CNV is typically gained as a tandem duplication, although PSC lines with four or five copies of this CNV have been reported^2,7^. The length of the duplicated region is also variable between different lines and ranges from 0.6Mb to 4Mb^2,7^. Nonetheless, the shared overlapping region in all of the reported variants contains a dosage-sensitive gene, *BCL2L1*, which was identified as the driver gene responsible for the key phenotypic features of variant PSC^5-7^. The altered behaviour of PSC harbouring the 20q11.21 CNV, coupled with the finding that the same CNV is a genomic hallmark of some cancers^8^, represents a potential impediment to the use of PSC in regenerative medicine applications and necessitates an understanding of the mechanisms governing the CNV appearance.

CNVs can arise as a consequence of DNA replication errors or during the process of DNA repair, with each of the implicated mechanisms of CNV formation yielding a different sequence profile within the resulting breakpoint junction^1^. For example, CNV formation can occur by the non-homologous end joining pathway when repair of DNA double strand breaks erroneously involves ligating the broken ends of different breaks instead of re-ligating the original site^9^. The editing of the broken ends prior to ligation is performed without the use of a homologous template and, consequently, the resulting breakpoint junctions in CNVs created by the non-homologous end joining typically contain random bases with no or little homology to the original sequence^10,11^. An alternative DNA repair mechanism implicated in CNV formation involves the non-allelic homologous recombination pathway, which drives the recombination of non-allelic genomic regions that share high sequence similarity, such as low copy repeats^1^. A defining feature of CNVs arising through this mechanism are long stretches of homology in the sequence flanking their breakpoints^12^. Finally, replication-based repair mechanisms of DNA repair, including fork stalling and template switching, and microhomology-mediated break-induced replication, can create CNVs by switching the nascent DNA strand from a stalled or collapsed replication fork to another fork in its vicinity, thereby giving rise to an insertion or a deletion of a DNA segment^13,14^. Importantly, invasion of an alternative replication fork requires a small region of homology with the complementary strand in order to prime the DNA synthesis. Therefore, CNVs formed by replication-based repair mechanisms are characterised by the presence of microhomology within their breakpoint sequence^14^.

Although the CNV genomic sequence holds essential clues as to the mechanisms governing its formation, this information is not attainable from conventionally employed techniques for CNV detection, such as the CGH arrays, Fluorescent In Situ Hybridisation or quantitative PCR^15^. By contrast, next generation sequencing technology can be used to reveal the CNV sequence at the nucleotide level, with increased or decreased numbers of mapped reads across genomic regions indicating the presence of genomic amplifications or deletions, respectively^16^. However, sequencing of the genome using short reads (<300bp) is ill-suited for CNV detection due to the mapping ambiguity of short reads, particularly in the presence of highly homologous or repetitive sequences^17^. Recently, the advent of long read sequencing technologies allowed reads to be uniquely mapped to the reference genome, thus facilitating a more effective CNV detection and identification of previously cryptic CNV breakpoints^18^.

Here, we applied long-range next generation sequencing to two human PSC lines that each harbour a 20q11.21 CNV, in order to delineate the CNV breakpoint sequences, the orientation of the amplified segments and the genomic context surrounding the CNV. The amplified segments were present in a head-to-tail orientation in both of the lines and their breakpoints contained sequences of microhomology, suggesting that the replication-based template switching mechanisms were implicated in their genesis. Moreover, we identified *Alu* repetitive sequences that intersect or flank the 20q11.21 CNV breakpoints. The presence of such repetitive elements may cause inherent instability to this area of the genome, making it a particular hotspot for CNV formation.

## Results

### Detection of human PSC lines with chromosome 20q11.21 CNV

By interphase FISH analysis, the human embryonic stem cell (ESC) line MShef7-A4, a subline of MShef7^19,20^, and the human induced pluripotent stem cell (iPSC) line NCRM1^21^ each exhibited a homogeneous population of cells with a tandem duplication or a triplication of the chromosome 20q11.21 region, respectively (**Supplementary Fig. 1**). To identify the approximate proximal and distal breakpoint position of the amplicon in each cell line (**Fig. 1**), we adapted our previously published qPCR-based method for assessment of copy number of target loci and we used it to assess the copy numbers of loci along the length of the q arm of chromosome 20^15,22^. In both cell lines, the proximal breakpoint was positioned between the centromere and the *DEFB115* gene (**Fig. 1**). In MShef7-A4, the distal breakpoint of the tandem duplication was located between the *TM9SF4* and *ASXL1* genes (**Fig. 1a, b**), whereas in NCRM1 the amplicon was smaller with the distal breakpoint positioned between the *TPX2* and *MYLK2* genes (**Fig. 1a, c**). In addition to identifying the putative breakpoints at 20q11.21, qPCR analysis revealed the presence of four copies of the amplicon in NCRM1, confirming the triplication of the chromosome 20q11.21 region in this line (**Fig. 1c**).

**Figure 1.**
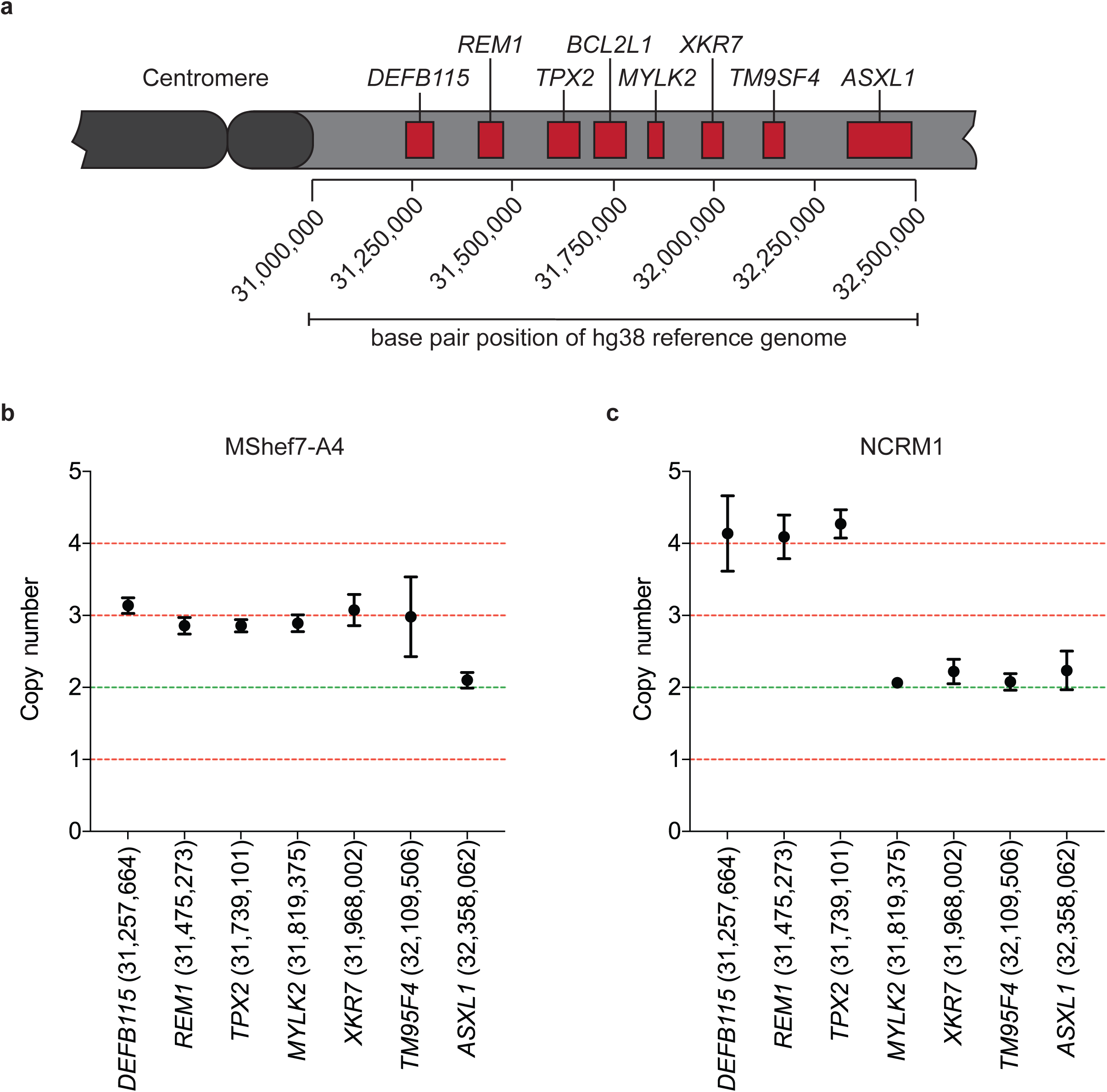
qPCR detection of distal breakpoint positions. **a**, A schematic showing the position and order of genes probed by qPCR along the chromosome 20q11.21. Primer sets were designed to target intronic regions of the genes displayed. **b**, Copy number values for the human ESC line MShef7-A4, determined by qPCR for loci along the length of chromosome 20q11.21. The primer location according to the hg38 reference genome are also displayed with the gene names along the X axis. **c**, The qPCR determined copy number for loci along the length of chromosome 20q11.21 in the NCRM1 human iPSC line. The copy number of four between *DEFB115* and *TPX2* indicates a triplication of this region.

### Nanopore sequencing reveals the chromosome 20q11.21 breakpoint in MShef7-A4

To identify the location of the breakpoints at a single nucleotide resolution in MShef7-A4 CNV and to determine the orientation of this tandem duplication, we performed whole-genome Oxford Nanopore sequencing on DNA extracted from the cells and aligned the sequencing reads to the hg38 human reference genome assembly^23^. The average read depth across chromosome 20 was 14.5 with a mean read length of 15.2 kb. We noted an increased sequencing read depth along the chromosome 20q11.21 relative to the rest of the chromosome (22.8 versus 14.5, respectively), indicative of a change in the copy number of this region (**Fig. 2a**)^24,25^. A distinct drop in read coverage was observed at position 32,273,600 bp of the chromosome 20 hg38 reference sequence (between *TPX2* and *MYLK2* genes), which we surmised was to be the distal breakpoint and was in agreement with the position we defined by qPCR (**Fig. 1a and 2a**). To represent reads which map to two discontinuous locations in the genome, mapping algorithms “soft-clip” reads to indicate that a portion of the read in question does not map to the same position as the remainder of the read. Soft-clipping of reads therefore provides evidence of structural variation, in our case, tandem duplication, as reads which span breakpoints map to disparate regions therefore triggering soft-clipping (**Supplementary Fig. 2**)^26,27^. Furthermore, the soft-clipped proportion of the sequencing read at the distal breakpoint can be used to infer the orientation of the tandem duplication. We reasoned that, if the soft-clipped DNA sequence at the distal breakpoint aligns to the reference genome between the centromere and *DEFB115* gene, then these two distantly positioned DNA sequences must have been fused in a head-to-tail orientation. However, if the soft-clipped portion of reads aligns to the distal breakpoint in an inverted orientation, the duplication has occurred in a head-to-head fashion. Therefore, we performed a BLAT pairwise sequence alignment of a contig formed from the unmapped portion of the soft-clipped reads to identify their genomic location^28^. The contig aligned with 92% identity to a (GGAAT)n microsatellite repeat in the pericentromeric region proximal of the *DEFB115* gene, confirming the head-to-tail orientation of the tandem duplication (**Fig. 2b, c**). This microsatellite is positioned at 31,051,509-31,107,036 bp on chromosome 20, and is flanked by two unmapped regions of the reference genome. We could not locate the proximal breakpoint to a single nucleotide position, which we inferred was due to the breakpoint being located in a currently unmapped region of the reference genome, potentially in one of the regions we observed flanking the microsatellite.

**Figure 2.**
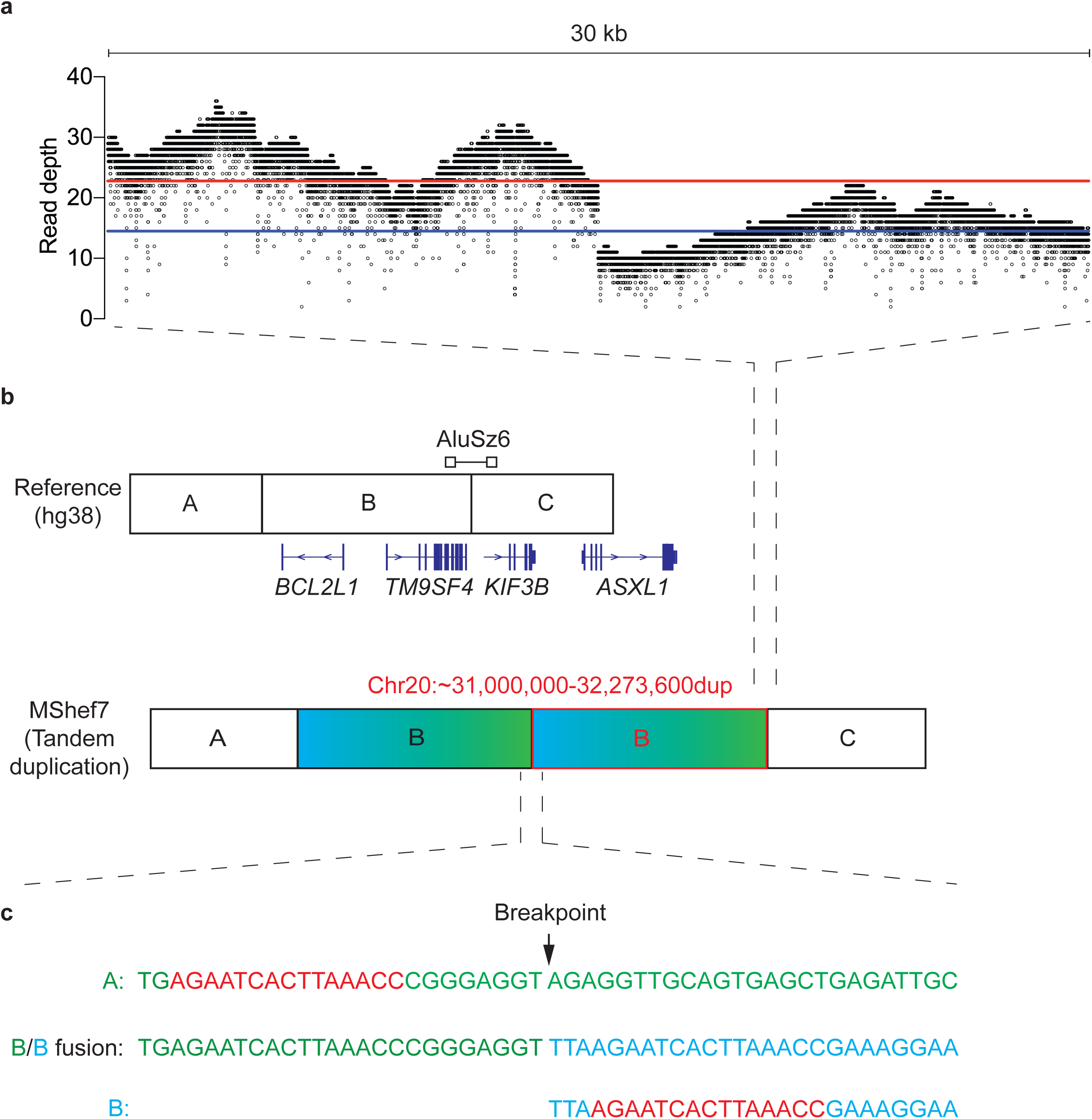
Breakpoint junction detection in MShef7-A4 using Nanopore sequencing. **a**, Sequencing read coverage of 30 kb spanning the breakpoint junction at 32,273,600 bp (chromosome 20q11.21) of the hg38 reference genome. Each dot indicates the read depth at a single base pair position. The red and blue lines indicate the mean read depth before and after the breakpoint position, respectively **b**, Schematic of the reference genome and the tandem duplication detected in MShef7-A4. Junction between genome segment A-B and B-C represents the proximal and distal breakpoint, respectively. The position of genes flanking and the location of the *AluSz6* in relation to the breakpoint are depicted. **c**, Reference sequence spanning the distal breakpoint (B – top, green), sequence of the breakpoint junction (B/B fusion– middle) and the contig sequence of the distal side of the proximal breakpoint (B – bottom, blue). The regions of microhomology that flank the proximal and distal breakpoint is highlighted (red).

To understand the mechanism of tandem duplication in MShef7-A4, we analysed the breakpoint sequences for signatures commonly observed in copy number variants. For the distal breakpoint, we analysed 500 bp of the reference genome sequence (hg38) surrounding the junction (**Fig. 2c**). As we were unable to locate the proximal breakpoint, we used the contig of the unmapped portions of the soft-clipped reads found at the distal breakpoint (**Fig. 2b, c**), which revealed a region of micro-homology (AGAATCACTTAAACC) that flanked both the proximal and distal breakpoint positions (**Fig. 2c**). By consulting the Dfam database of transposable elements, we observed that the distal region of microhomology lies within an *AluSz6* retrotransposon that spans the distal breakpoint^29^. These results suggest a role of microhomology in the mutational mechanism of the tandem amplification of chromosome 20 in the MShef7-A4 cell line.

### Break point mapping of a chromosome 20q11.21 tandem triplication

We used the same sequencing approach to identify and analyse the breakpoints in the human iPSC cell line, NCRM1, which contains a tandem triplication in the 20q11.21 region (**Supplementary Fig. 2**). Our Nanopore sequencing returned an average read length of 19.9 kb at a mean depth of 20.3 across chromosome 20. Consistent with our qPCR analysis, long-read sequencing identified a sole distal breakpoint at position 31,813,288 bp between the *TPX2* and *MYLK2* genes. This confirmed that both amplicon copies in NCRM1 have the same distal breakpoint position. The increased read depth associated with copy number variants was greater in NCRM1 (43.9) when compared with MShef7-A4, consistent with the triplication indicated by our PCR and FISH analyses (**Fig. 3a**). To identify the proximal breakpoint position, we performed a BLAT pairwise sequence alignment on the unmapped portions of the soft-clipped reads. Our soft-clipped sequence aligned with the reference genome at position 31,059,954 bp, within the same microsatellite that was putatively identified as the proximal breakpoint region in MShef7-A4 (**Fig. 3b, c**). These data confirm that the tandem triplication of chromosome 20q11.21 in NCRM1 has occurred in a head-to-tail orientation, and that each amplicon was of equal length and contained the same breakpoint positions. Furthermore, we observed a common microsatellite sequence at the proximal breakpoint in both cell lines, and thus, its involvement could be complicit in the tandem amplifications that commonly occur associated with chromosome 20q11.21.

**Figure 3.**
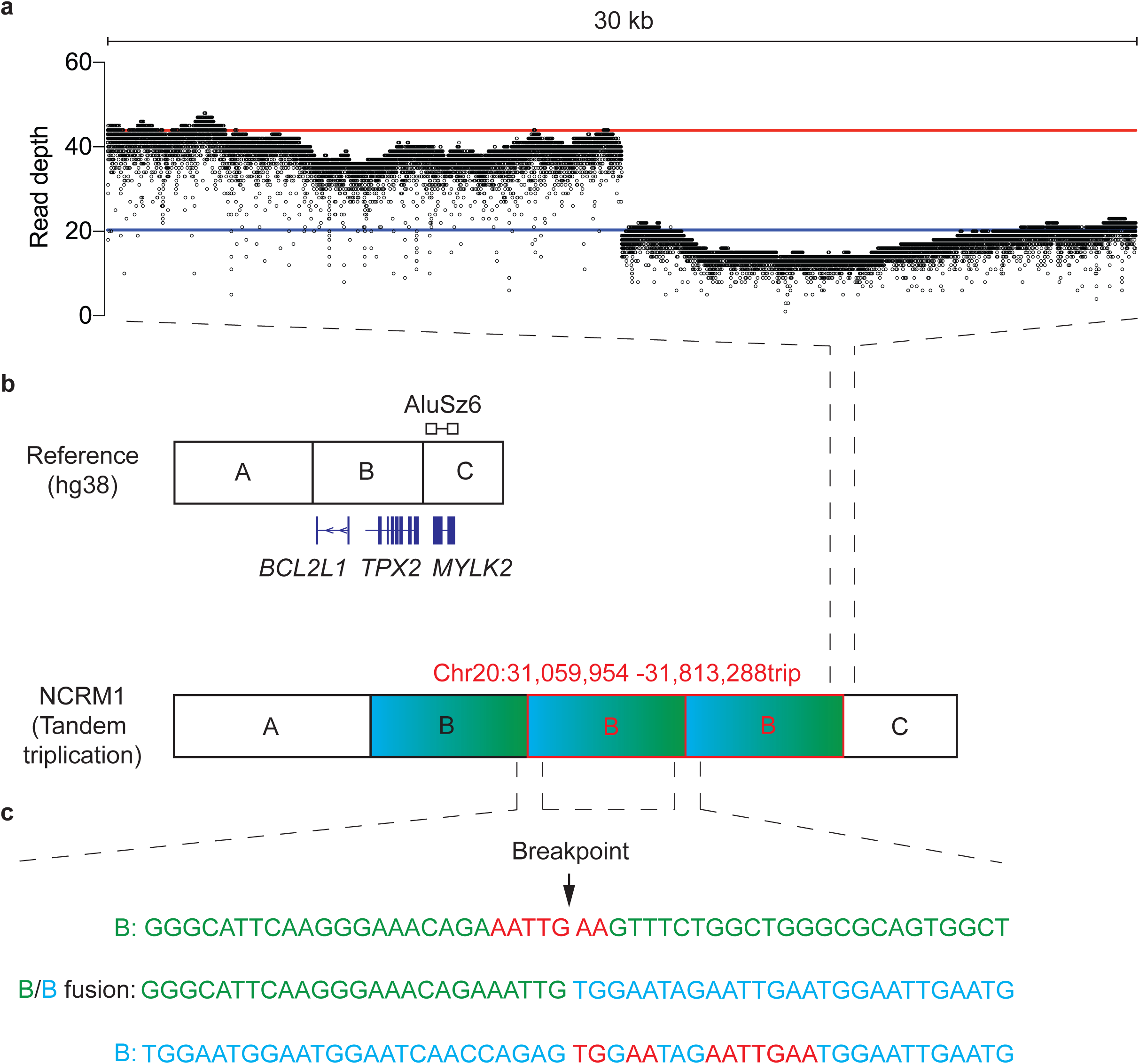
Breakpoint position of the tandem triplication in NCRM1. **a**, Read coverage of 30 kb surrounding the breakpoint junction 31,813,288 bp (chromosome 20q11.21) of the hg38 reference genome. The mean read depth before and after the breakpoint is shown (red line and blue line, respectively). **b**, Schematic depicting the reference genome and the NCRM1 tandem triplication. The distal breakpoint lies between the junction of B-C and the proximal breakpoint is located on the boundary of the A-B segments. The genes flanking the breakpoint, as determined by qPCR are depicted. The position of the *AluSz6* identified from the Dfam database is represented above the reference sequence schematic. The exact nucleotide position of the proximal and distal breakpoint is written above the schematic of the tandem triplication. **c**, Reference sequence spanning the distal breakpoint (B – top, green), the proximal breakpoint (B – bottom, blue) and the combined amplification breakpoint junction (B/B fusion – middle). The region of microhomology that flanks each of the breakpoints is highlighted (red).

To infer the mechanism involved in the tandem triplication of chromosome 20q11.21 in NCRM1, we interrogated the reference sequence at both the proximal and distal breakpoint positions. We identified multiple regions of micro-homology (TGAA and AATTGAA) that flanked both sides of the fusion junction (**Fig. 3c**). Furthermore, we consulted the Dfam database of transposable elements and identified an *AluSz6* element that was situated 9 bp downstream of the distal breakpoint (**Fig. 3b, c**). As we were unable to find an *Alu* element at the proximal breakpoint itself, it is unlikely the tandem duplication and triplication in MShef7-A4 and NCRM1, respectively, have arisen through a mechanism of *Alu-Alu* recombination. Instead, we propose that the *Alu* elements are sites of chromosome fragility, due to replication blockage^30-34^. Repair of stalled and collapsed forks would then proceed through replication fork switching to complementary sites of microhomology, and strand invasion upstream on the same or a homologous chromosome would generate a tandem amplification (**Fig. 4**).

**Figure 4.**
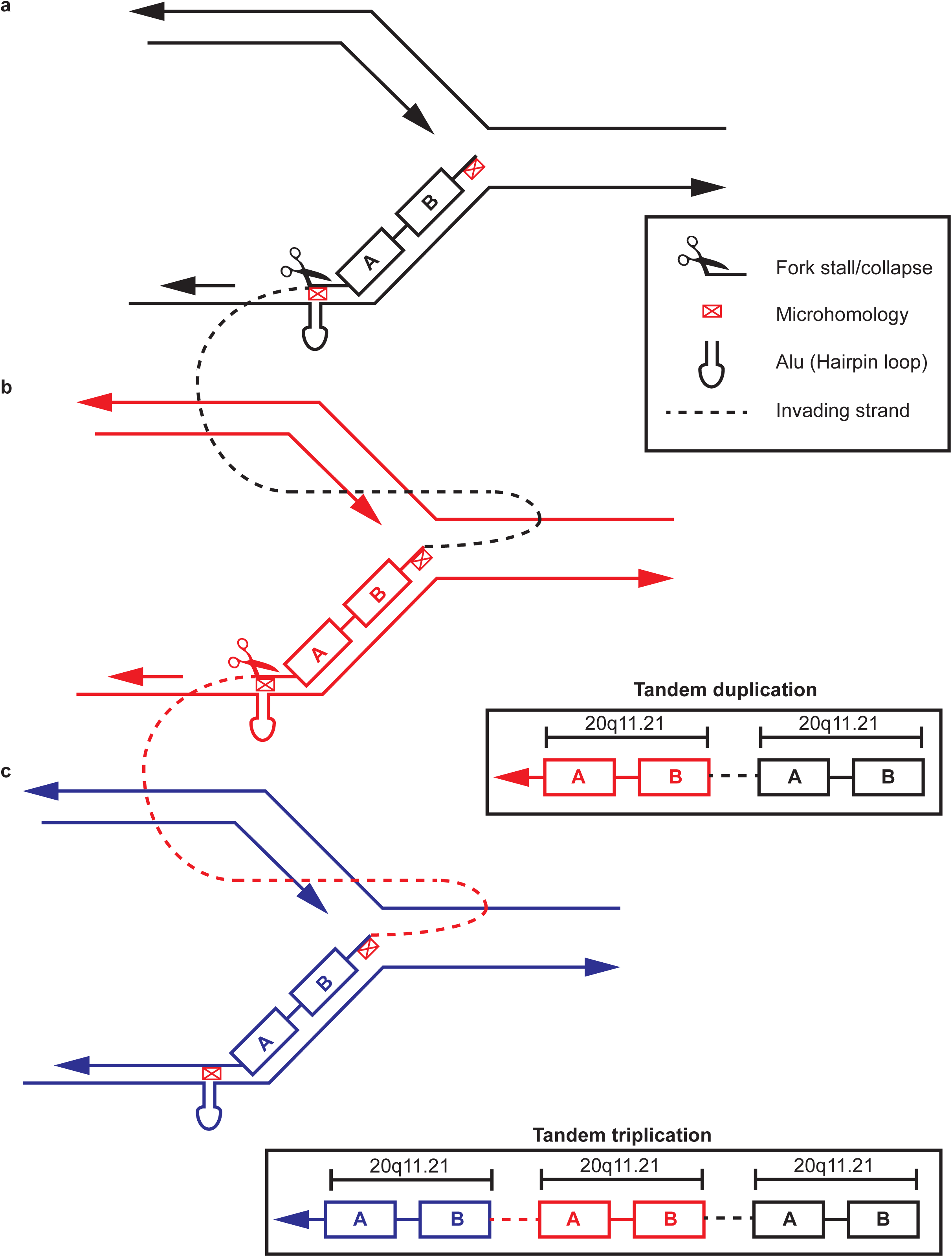
Replication template switching is responsible for tandem amplification in human PSC. **a**, Replication fork stalling is promoted by *Alu* sequences that form hairpin loops. **b**, Replication fork repair by fork stalling and template switching and/or microhomology mediated break induced replication is initiated by strand invasion at a site of microhomology in the pericentromeric microsatellite on the sister chromatid. Replication proceeds, duplicating 20q11.21. **c**, An additional round of strand invasion and re-synthesis occurs of the other parent homolog in examples of tandem triplication.

## Discussion

The experiments reported here have revealed the breakpoints of tandem amplifications of chromosome 20q11.21 in human PSC. The distal breakpoints were all found to be located in, or close to *Alu* sequences. The proximal breakpoints were located in a pericentromeric microsatellite region close to 31 Mb on chromosome 20. In the case of NCRM1, each amplicon of the tandem triplication was of equal length with the same breakpoint positions. A detailed characterisation of the breakpoints at a single nucleotide level revealed short microhomologies that flank or overlap both the proximal and distal breakpoints. These breakpoint characteristics are like scars left by the repair mechanism that operated on the DNA lesion.

Although rare, breakpoint microhomology of between 1-4 bp long is occasionally observed with CNV formed by non-homologous end joining (NHEJ)^35,36^. As the microhomology at the breakpoints of our lines was larger than 7 bp we excluded classical NHEJ as the mechanism of tandem amplification. However, alternative forms of end-joining such as microhomology mediated end joining do utilize larger spans of homology or microhomology^37-42^. These mechanisms differ from classical NHEJ, as they do not perform blunt-end ligation and instead utilise end-resection at DNA breaks to reveal overlapping micro-homologous single stranded DNA required for annealing^43^. We eliminated alternative end-joining from the potential mutagenic mechanisms, as the microhomology in both MShef7 and NCRM1 was intact and tandem amplifications are not readily explained by this mechanism ^44^.

The tandem amplifications in MShef7 and NCRM1 had breakpoints devoid of large regions of sequence homology, which ruled out mechanisms involving homologous recombination such as non-allelic homologous recombination ^45^. However, the presence of an *AluSz6* element at the distal breakpoints in both cell lines led us to consider *Alu-Alu*-mediated non-allelic homologous recombination mechanism. For *Alu-Alu*-mediated non-allelic homologous recombination to take place it would require a second *Alu* element at the proximal breakpoint with high sequence identity with the distal *Alu*^46^. We found no evidence of a second *Alu* at the proximal breakpoint in either of our cell lines. Despite this, the presence of *AluSz6* at distal breakpoints in both cell lines suggests that it might play a role in the initiation of tandem amplifications, rather than in the mechanism of mutation itself. Inverted repeats, such as *Alu* elements, form hairpin loop secondary structures that can impede replication, leading to fork stalling and collapse, particularly under conditions of replication stress^30-34,47-49^. It is perhaps no coincidence then, that this mechanism of mutagenesis is associated with high levels of replication stress, which is a characteristic of human PSC during *in vitro* culture^50-52^.

The breakpoint signatures of the tandem amplifications characterised in MShef7-A4 and NCRM1 are consistent with the replication template switching mechanisms, fork stalling and template switching and microhomology mediated break induced replication, which are initiated by replication fork stalling and collapse, respectively^13,14^. In the case of fork stalling and template switching, the lagging strand at the stalled fork disengages and invades another replication fork at a region of microhomology. Microhomology mediated break induced replication is similar to fork stalling and template switching, although following a collapsed fork the 5’ end of the DNA break is resected to generate a 3’ single-stranded overhang that then invades a template region with microhomology before replication is reinitiated. If the template is upstream on the same chromosome or a homologous chromosome, a tandem amplification would result (**Fig. 4a, b**)^13,14,45,53^. Furthermore, the role of microhomology mediated break induced replication and fork stalling and template switching in the formation of tandem triplications has been discussed^14,54-56^. Should replication fork collapse lead to sister chromatid strand invasion at an upstream region of microhomology, replication of the amplified segment will proceed. This could then be followed by a second round of template switching and strand invasion at the same region of microhomology, although this time into the other parental homolog with replication proceeding to the distal end of the chromosome, resulting in a tandem triplication (**Fig. 4a-c**)

In summary, we provide evidence from breakpoint junctions that implicate replication-based repair by fork stalling and template switching and microhomology mediated break induced replication as the mutational mechanism driving tandem duplication in human PSC. We argue that constitutive replication stress observed during the *in vitro* culture of human PSC could be driving replication fork stalling and collapse at *Alu* elements that initiates these mutations. This report provides new insight into the mechanisms of mutation in human PSC. The recurrent nature of genetic change in human PSC is considered non-random due to the selection of advantageous mutations. However, it was recently reported that mutations in human PSC occur with higher frequency in non-genic regions^57^. The data presented here complements these findings and suggests that mutation itself may be non-random but may be enriched at certain sites that can be characterised by the genomic architecture. By defining these regions, it may be possible to safeguard the genome stability of human PSC for their use in cell-based regenerative medicine.

## Methods

### Human pluripotent stem cell culture

The MShef7^19,20^ (hPSCreg: https://hpscreg.eu/cell-line/UOSe012-A) human ESC line was derived at the University of Sheffield Centre for Stem Cell Biology under the HFEA licence R0115-8A (centre 0191) and HTA licence 22510. A mosaic sub-population of chromosome 20 variant cells was detected in a culture of MShef7, which was sub-cloned using single cell deposition by FACS. The NCRM1^21^ (hPSCreg: https://hpscreg.eu/cell-line/CRMi003-A) human iPSC line was acquired from RUCDR Infinite Biologics and was originally derived by reprogramming umbilical cord blood CD34+ cells using a non-integrating episomal vector. Both cell lines were maintained in culture vessels coated with a matrix of Vitronectin human recombinant protein (ThermoFisher Scientific, A14700) and batch fed daily with mTeSR (STEMCELL Technologies, 85850). Once the cells had reached confluency, they were passaged using ReLeSR (STEMCELL Technologies, 05873) according to manufacturer’s guidelines.

### qPCR breakpoint determination

DNA was extracted from cell pellets using the DNeasy Blood and Tissue kit (Qiagen, 69504). DNA quantity and quality were measured using a NanoPhotometer (Implen). 1μg of DNA was digested with 10 units of FastDigest EcoRI enzyme (Thermo Fisher Scientific, FD0275) in FastDigest buffer (Thermo Fisher Scientific, FD0275) for 5 minutes at 37°C, followed by deactivation of the enzyme by incubating at 80°C for 5 minutes. qPCR was performed as previously described^15,22^, using the adapted protocol^22^ whereby primer sets were designed along the length of the q arm of chromosome 20 (**Table 1**) to allow an estimate of the amplicon length. A 10µl PCR reaction contained TaqMan Fast Universal PCR mastermix (ThermoFisher Scientific, 4366072), 0.1 μM Universal probe library hydrolysis probe, 0.1 μM each of the forward and reverse primers (**Table 1**) and either 20ng of EcoRI-digested DNA or water only (no template control). The PCR reactions were run on the QuantStudio 12K Flex Real-Time PCR System using the following profile: 50°C for 2 minutes, 95°C for 10 minutes, and 40 cycles of 95°C for 15 seconds and 60°C for 1 minute. The copy number was determined by first subtracting the average Cq values from the test sample 20q loci from the reference loci (Chromosome 4p) to obtain a dCq value. The dCq for the calibrator sample at the same loci was then calculated in the same way and the test sample dCq and calibrator sample dCq were subtracted from one another to give ddCq. The relative quantity was calculated as 2^-ddCq^. Finally, to obtain the copy number, the relative quantity value was multiplied by 2.

**Table 1.**
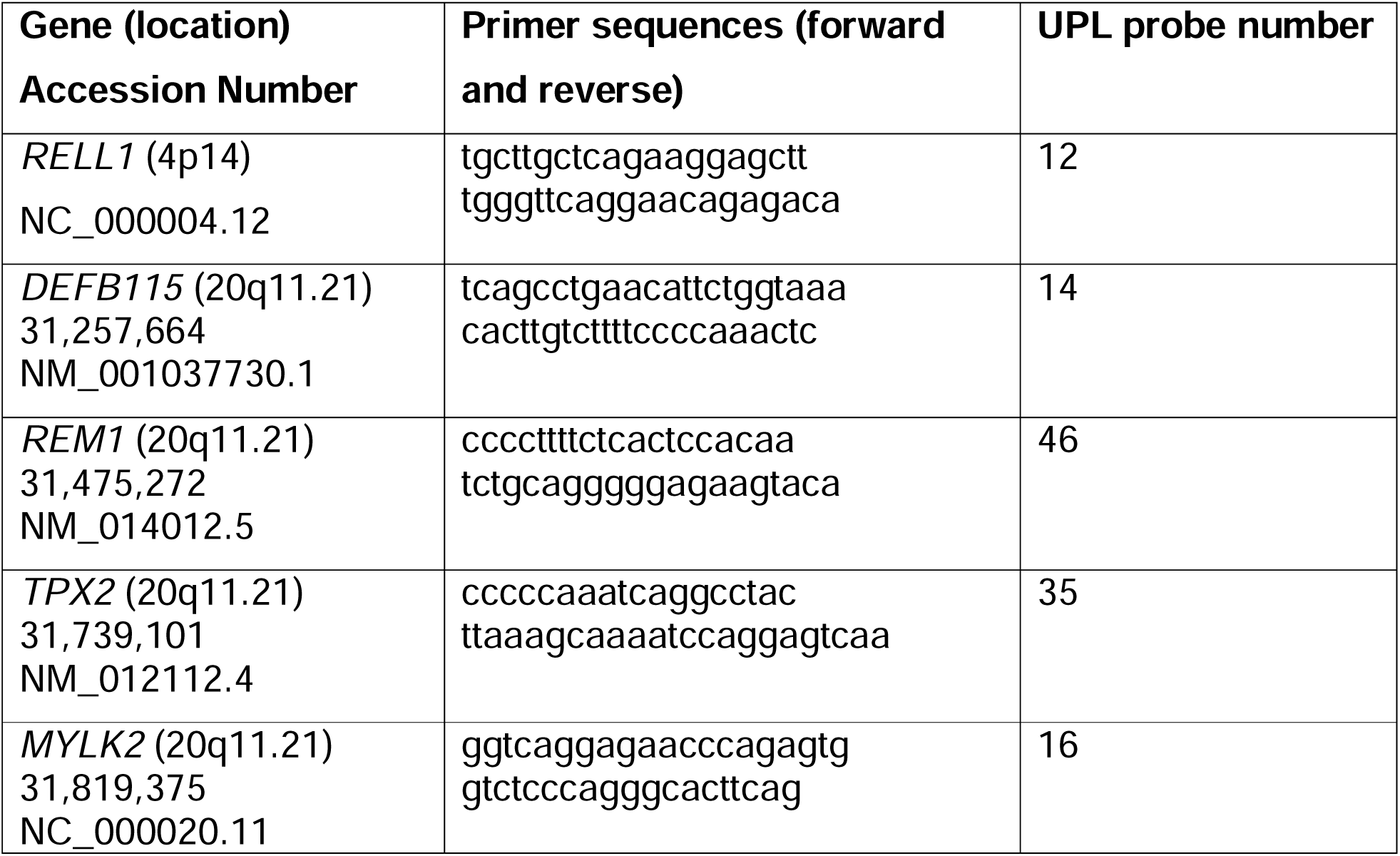
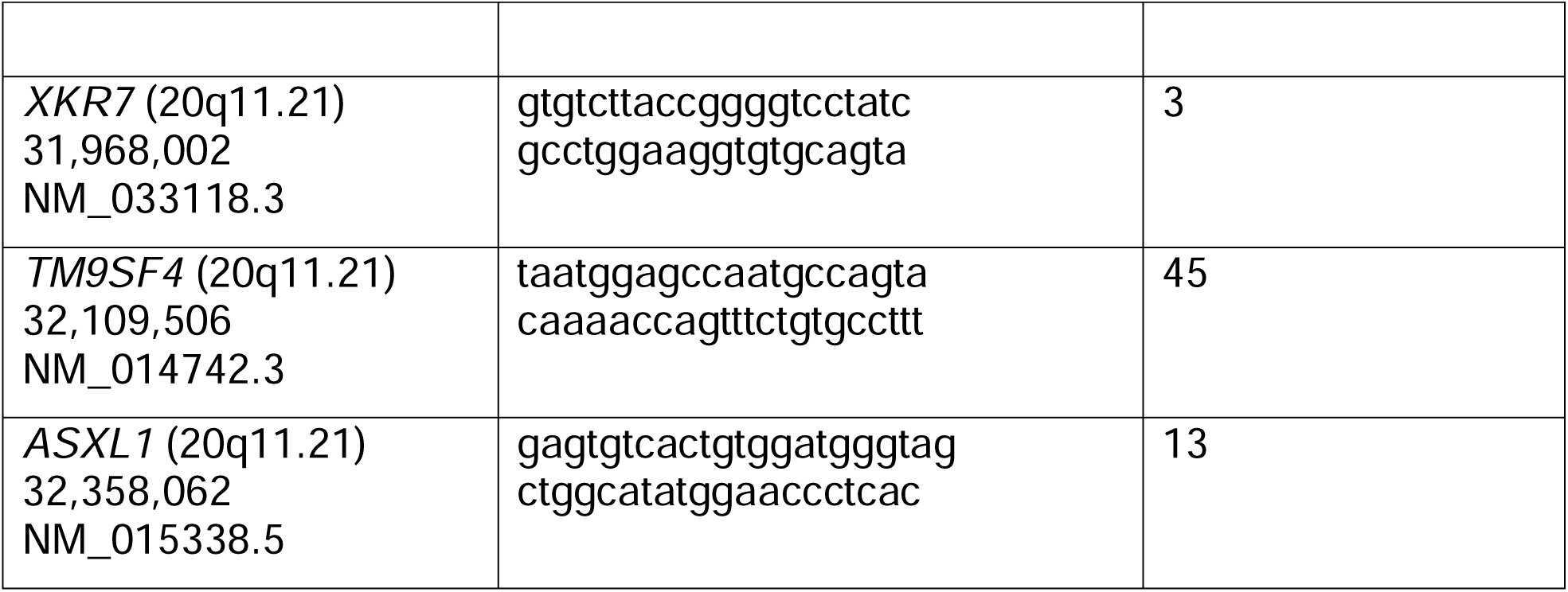
qPCR breakpoint detection primer sets and probes ^22^.

### Fluorescence *in situ* hybridisation (FISH) for the detection of chromosomal variants

Human PSC were detached from culture flasks by incubating with TrypLE Express Enzyme (Fisher Scientific, 11528856) for 3 minutes at 37°C. The cells were collected in DMEM/F12 basal media (D6421, Sigma Aldrich) and centrifuged at 270 g for 8 minutes. To the cell pellet, 1 mL of pre-warmed 37°C 0.0375 M potassium chloride was added. The cells were then centrifuged at 270 g for 8 minutes, before fixing the cells by adding 2 mL fixative (3 parts methanol: 1 part acetic acid, v/v), in a drop-wise manner under constant agitation. FISH detection of chromosomal variants was performed by Sheffield Diagnostics Genetic Service. Analysis was performed on 100 interphase nuclei per sample that had been probed with D20S108 (BCL2L1) probe.

### DNA extraction for sequencing

DNA was extracted from cell pellets using the DNeasy Blood and Tissue kit (Qiagen, 69504). DNA quantity and quality were measured using a NanoPhotometer (Implen).

### DNA sequencing

DNA library preparation was performed using the ligation (Oxford Nanopore Technologies, SQK-LSK108) or Rapid sequencing kits (Oxford Nanopore Technologies, SQK-RAD004) according to the manufacturer’s Genomic DNA by Ligation or Rapid Sequencing protocols, respectively. The whole genome libraries were sequenced using the Oxford Nanopore MinION or GridION sequencers with the R9.4.1 flow cell (Oxford Nanopore Technologies, FLO-MIN106D) following the manufacturer’s instructions. Each flow cell yielded ∼5 Gb of data.

### Data processing

Data exported as FASTQ files were mapped to the chromosome 20 hg38 reference sequence using minimap2 sequence aligner (version 2-2.15)^58^. File management, merging, sorting and indexing was performed using Sambamba (version 0.6.6) and Samtools (version 1.9)^59,60^. Breakpoint regions were inspected manually using IGV genomic viewer^61^ and the breakpoint location was identified based on read depth and soft-clipped sequence analysis. Briefly, the aligned and sorted .bam files were opened using IGV genomic viewer with soft-clipped bases enabled. The distal breakpoint region identified by qPCR was inspected and the breakpoint at the single nucleotide level was located by identifying a region of reduced read depth with soft-clipped reads that spanned the point of reduced read coverage (**Figure S2A, B**). To identify the proximal breakpoint, the soft-clipped proportion of the sequencing reads at the distal breakpoint were queried using BLAT sequence alignment to identify the sequence matches in the human reference genome with high similarity.

## Acknowledgements

The authors would like to thank Matthew Parker, Emily Chambers and Mark Dunning of the Sheffield Bioinformatics Core, The University of Sheffield for assistance and advice with performing the data processing. This work was partly funded by the European Union’s Horizon 2020 research and innovation program under grant agreement No. 668724 and partly by the UK Regenerative Medicine Platform, MRC reference MR/R015724/1. The Wellcome Sanger Institute is grateful for the Wellcome Trust general core grant number 206194.

## Author Contribution

PWA and IB oversaw the project. JAH, PWA and IB devised the experimental design. JAH performed the cell culture, DNA extraction, qPCR and data processing. Additional help for data processing was provided by the Sheffield Bioinformatics Core. DB performed interphase FISH detection of chromosome 20 amplification. KJ, MAQ, KO, EB and JS performed the Nanopore library preparation and whole genome sequencing. The manuscript was drafted by JAH, PWA and IB.

