## Supplementary Fig. for "Template switching mechanism drives the tandem amplification of chromosome 20q11.21 in human pluripotent stem cells"

**Supplementary Information**

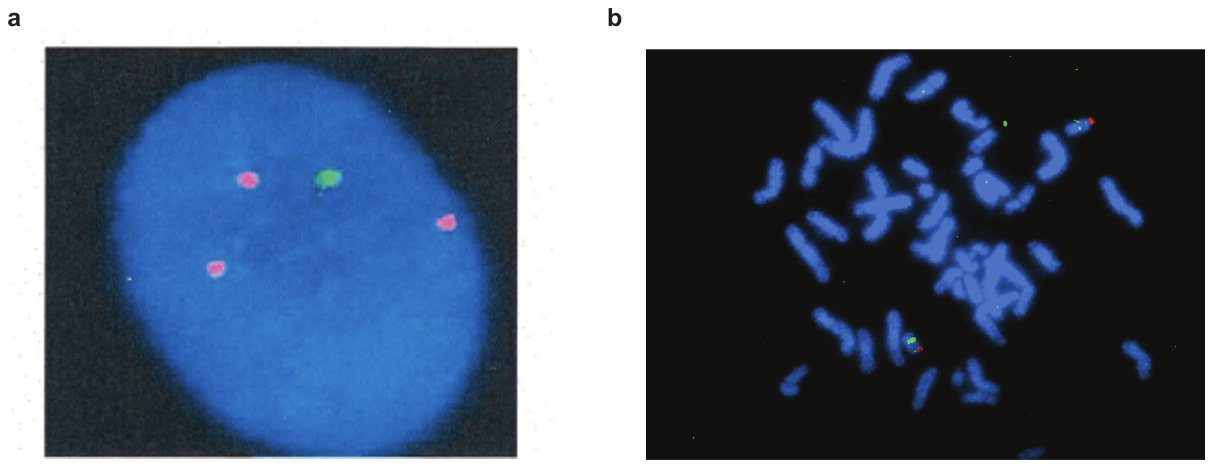

**Supplementary Figure 1 | Interphase and metaphase FISH detection of chromosome 20 tandem amplification.** Representative image of interphase FISH performed on MShef7-A4 (**a**) and metaphase FISH performed on NCRM1 (**b**) hybridised with D20S108 q11.21 probe (red) and chromosome 20 centromere probe (green), DNA was co-stained with DAPI (Blue).

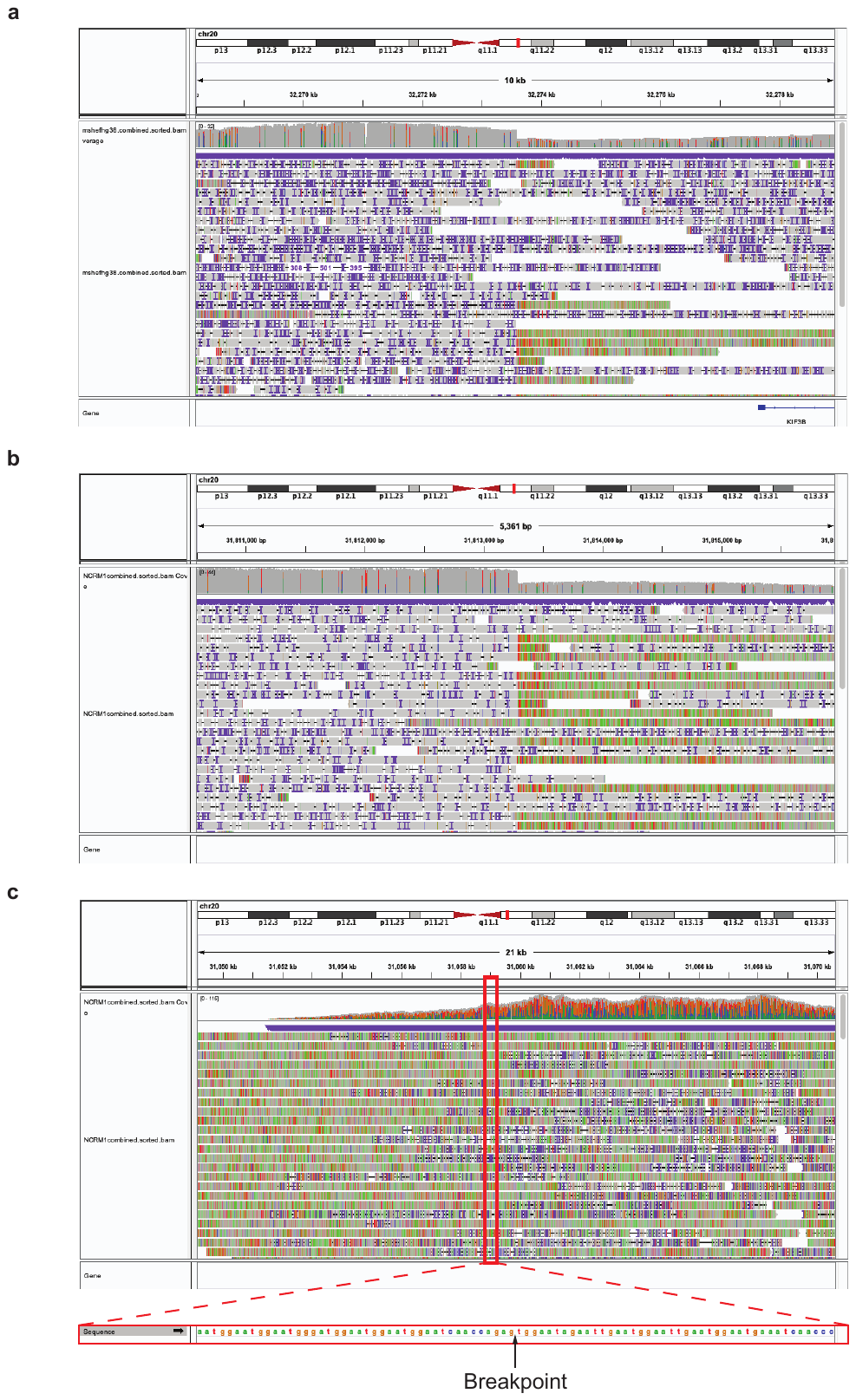

**Supplementary Figure 2 | Breakpoints identification in IGV genomics viewer. a,** Screenshot of the distal breakpoint position in MShef7-A4 (32,273,600 bp). Read depth shows a distinctive drop in coverage at the centre of the image that coincides with soft clipped reads that flank downstream of this position. **b,** IGV image of the distal breakpoint (31,813,288 bp) in NCRM1. The breakpoint was identified from read depth change and soft-clipped read identification. **c,** The soft-clipped reads from the distal breakpoint mapped to 31,059,954 bp. Inset, reference sequence surrounding the breakpoint position.
